# Shroom3-Rock interaction and profibrotic function: Resolving mechanism of an intronic CKD risk allele

**DOI:** 10.1101/2024.11.22.624409

**Authors:** Anand Reghuvaran, Ashwani Kumar, Qisheng Lin, Nallakandi Rajeevan, Zeguo Sun, Hongmei Shi, Gabriel Barsotti, EM Tanvir, John Pell, Sudhir Perincheri, Chengguo Wei, Marina Planoutene, Anne Eichmann, Valeria Mas, Weijia Zhang, Bhaskar Das, Lloyd Cantley, Leyuan Xu, Cijiang John He, Madhav C Menon

## Abstract

Common intronic enhancer SNPs in Shroom3 associate with CKD in GWAS, although there is paucity of detailed mechanism. Previously, we reported a role for Shroom3 in mediating crosstalk between TGFβ1- & Wnt/Ctnnb1 pathways promoting renal fibrosis (TIF). However, beneficial roles for Shroom3 in proteinuria have also been reported suggesting pleiotropic effects. Here we focused on identifying the specific profibrotic Shroom3 motif. Given known therapeutic roles for Rho-kinase inhibitors in experimental CKD, and the established interaction between Shroom3 and Rock via its ASD2 domain, we hypothesized that Shroom3-mediated ROCK activation played a crucial role in its profibrotic function in high expressors. To test this hypothesis, we developed transgenic mice and cell lines that inducibly overexpressed wild-type- (WT-Sh3) or ASD2-domain deletion- Shroom3 (ASD2Δ-Sh3). Prior scRNAseq data showed that during TIF, Shroom3 and Rock co-expression occurred in injured tubular cells and fibroblasts, highlighting cell-types where this mechanism could be involved. Using HEK293T cells, we first confirmed absent ROCK binding and inhibited TGFβ1-signaling with ASD2Δ-Sh3-overexpression vs WT-Sh3. In mIMCD cells, ASD2Δ-Sh3 overexpression, reduced Rock activation (phospho-MYPT1), pro-fibrotic and pro-inflammatory transcripts vs WT-Sh3. Fibroblast proliferation (3T3) was also reduced with ASD2Δ-Sh3. *In vivo*, we studied ureteric obstruction (UUO) and Aristolochic nephropathy (AAN) as TIF models. In AAN, inducible global-, or Pan-tubular specific-, WTSh3-overexpression showed increased azotemia, and TIF vs ASD2Δ-Sh3 mice. WT-Sh3 mice consistently showed significant enrichment of Rho-GTPase, TGFβ1- and Wnt/CtnnB1- signaling in kidney transcriptome, paralleling Shroom3-coexpressed genes in tubulo-interstitial transcriptomes from human CKD. In UUO, again WT-Sh3 mice recapitulated increased fibrosis vs ASD2Δ-Sh3. Importantly, ASD2Δ-Sh3 did not develop albuminuria vs WT-Sh3, while mutating a disparate Fyn-binding Shroom3 motif induced albuminuria in mice, suggesting motif-specific roles for Shroom3 in the kidney. Hence, our data show a critical role for the Rock-binding, ASD2-domain in mediating TIF in milieu of Shroom3 excess, with relevance to human CKD.

## Introduction

In native and allograft kidneys, CKD is characterized by progressive renal interstitial fibrosis and tubular atrophy, reflecting accruing damage. In humans, while GWAS have identified candidate CKD susceptibility loci(1, 2), the detailed mechanistic basis of SNP-variant associations with renal histology/function in CKD remain limited (3). This has hindered translation of therapeutics based on GWAS. Specifically, targeting fibrosis mechanisms in the native or allograft CKD context have not successfully translated to the bedside in clinical trials, in spite of promising preclinical data(4). As a result, a significant percentage of patients with susceptibility loci for CKD or allograft injury will eventually progress to ESRD requiring dialysis or transplantation(5, 6).

Within the Shroom3 gene locus, multiple <u>intronic SNPs including rs17319721 have associated with CKD in GWAS</u>(1, 2, 7). We and others identified that the A-allele at rs17319721 has TCF7L2-dependent enhancer function on *Shroom3* in renal epithelial cells(8, 9). In renal allograft biopsies, enhanced *SHROOM3* expression with the donor A-allele, associated with increased TGFβ1- & Wnt/Ctnnb1 pathways promoting renal fibrosis (TIF)(8) and reduced eGFR by 1-year post-transplant(10). We showed that SHROOM3 overexpression in tubular cells increases profibrotic marker production in response to TGF-β1, and inducible global or renal-tubular specific *Shroom3* knockdown in mice, significantly inhibited TIF(8). These data suggested a pathogenetic role for SHROOM3 in TIF, and potential for testing the role of SHROOM3-antagonism as a therapeutic strategy for TIF in CKD in A-allele kidneys. However, subsequent data <u>from our/other groups showed a protective role for Shroom3 in glomerular development</u>(11,12),(13)<u>, and indicated the development of proteinuria when Shroom3 was antagonized in adult podocytes</u>(14, 15).Together, these data suggest an adverse association for rs17319721 and elevated *Shroom3* with renal TIF in CKD, but also cautioned that globally antagonizing Shroom3 would incur the risk of proteinuria.

Shroom3 belongs to the *SHROOM* family of proteins(16) containing a PDZ domain, two Apx/Shrm-specific domains called ASD1 and ASD2(17). The ASD1 domain directly binds F-Actin (18) while the ASD2 domain binds and activates Rho-associated coiled-coil kinases (ROCKs)(19). We reported a previously unidentified -PxxP- domain located between PDZ and ASD1-domains, which was critical for Fyn-binding and activation in podocytes, and that adult mice with Shroom3 knockdown phenocopied Fyn-KO mice, with inhibited Nphs1 phosphorylation and Actin cytoskeletal organization demonstrating that this motif was crucial in adult podocyte homeostasis(14). Subsequently we reported that Shroom3 knockdown in glomerular podocytes induced a minimal change-like injury, due to Fyn inactivation with collateral activation of AMPK(15), and inhibition of AMPK in this context induced FSGS. Notably, impaired Fyn activation in podocytes has also been specifically associated with human MCD(20). These suggested that the Fyn-binding motif of Shroom3 mediated its antiproteinuric effects in adult animals, distinct from the developmental role in glomerulogenesis reported before(11, 12).

In this work, we performed a series of experiments which focused on identifying and specifically targeting the profibrotic motif of Shroom3. We first developed deletional variants of each of the consensus domains of Shroom3 and screened them invitro in multiple cell lines identifying a key role for the Rock-binding ASD2-domain in profibrotic signaling facilitating TGFβ1 and Wnt/Ctnnb1 pathways. These data were consistent with prior antifibrotic roles for Rock-inhibitor, Hydroxy-Fasudil(HF)(21–23). To study the *in vivo* role played by Shroom3 motifs in the development of TIF, we then developed novel inducible, cell-specific overexpression mice to induce either intact Shroom3 excess (analogous to the risk-allele carriage with high expressor kidneys) or overexpression of a variant of Shroom3 without the ASD2-motif. We compared these in multiple TIF models and demonstrate that the ASD2-motif is the profibrotic motif during TIF. Hence our work identifies underlying pathophysiologic mechanisms associating Shroom3 SNPs with CKD and allograft injury and is proof-of-principle for designing Shroom3 targeted precision therapeutics.

## Results

### 1. ASD2Δ-Sh3-overexpression reduces ROCK activation impairing pro-inflammatory and profibrotic signaling in tubular cells & fibroblasts vs WT-Sh3

Using published ScRNAseq data from murine kidneys, we first investigated cell-type specific expression of Shroom3 from TIF models (24, 25) [Fig 1A]. We focused on cells that co-expressed *Shroom3* with *Rock1* and/or *Rock2* during TIF. From UUO and post-ischemia-reperfusion (I-R) TIF we identified that injured PT cells, distal nephron epithelial cells (collecting duct/connecting tubule/intercalated cells) and fibroblasts/myofibroblasts show higher *Shroom3, Rock1, Rock2* expression. In a second dataset of post-IR TIF again, injured PT cells especially in the TIF stage, showed higher *Shroom3* & *Rock* expression[Fig S1A](26). Podocytes show high expression of Shroom3 at rest and injury but are not expected to play key roles in these TIF models after proximal tubular cell injury. Hence, we focused on tubular and fibroblast cell lines in subsequent mechanistic experiments.

**Figure 1:**
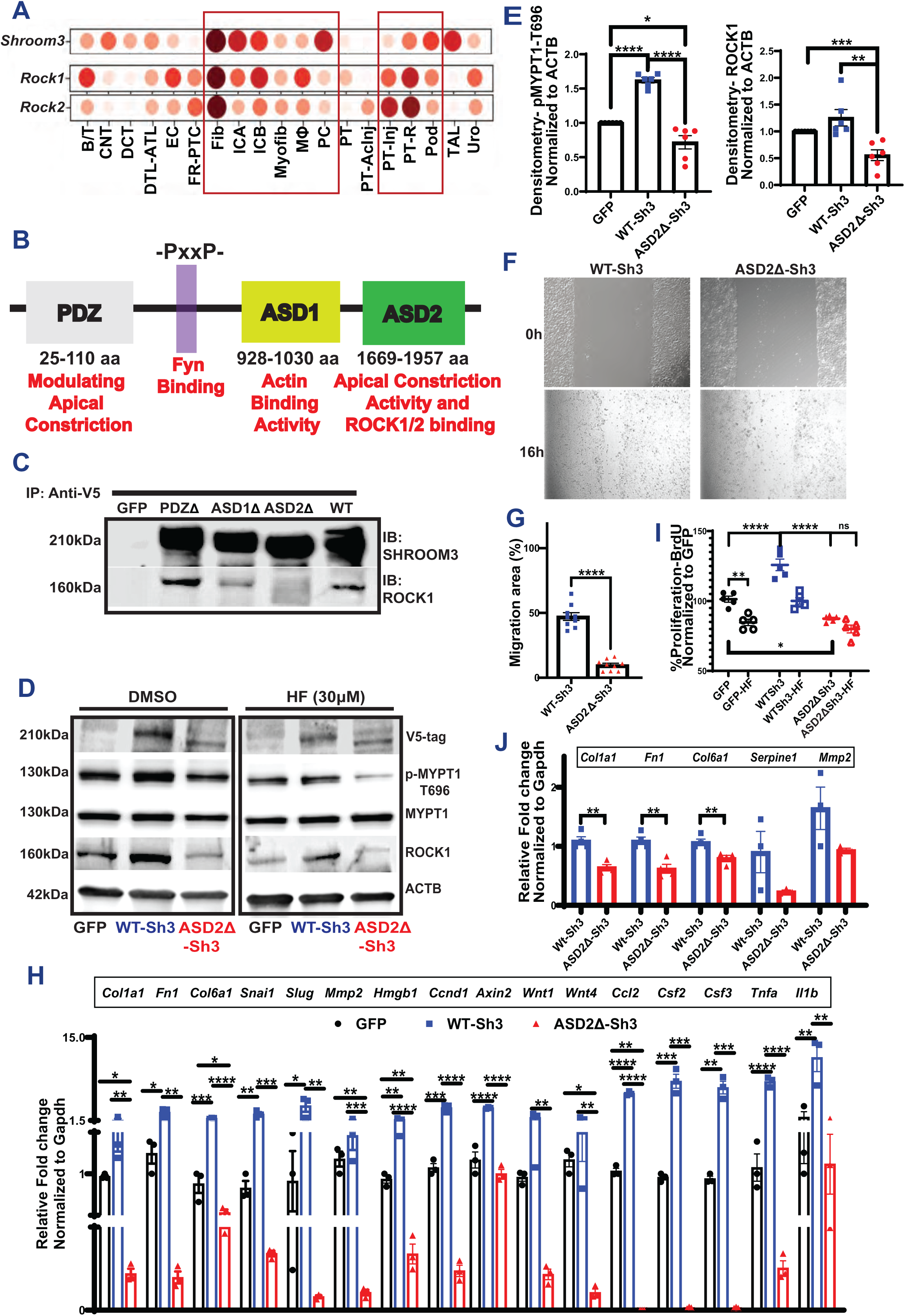
(**A**) Dot plot representing *Shroom3, Rock1, Rock2* in multiple cell types identified in Injured murine kidneys (https://humphreyslab.com/SingleCell/) (**B**) The conserved domains of Shroom3 are shown (PDZ, ASD1, ASD2 domains and the Fyn-binding site). (**C**) Representative immunoblots of SHROOM3 and ROCK1 from proteins immunoprecipitated from HEK293T cells overexpressing PDZ, ASD1 and ASD2 using V5 beads. (**D**) Representative immunoblots of lysates from Dmso/HF-treated mIMCD cells overexpressing GFP, WT-Sh3 and ASD2Δ-Sh3 for V5 (Shroom3), phospho-MYPT, total MYPT, ROCK1 and βACTIN. (**E**) Dot plots of relative quantification of phospho-MYPT and ROCK1. (**F**) Phase contrast micrographs of wound closure in mIMCD cells expressing WT-Sh3/ ASD2Δ-Sh3 and the (**G**) Bar graph representing the percent migrated area. (**H**) Bar graph representing the relative mRNA expression of markers of fibrosis, Wnt signals and pro-inflammatory chemokines in mIMCD cells (WT-Sh3- vs ASD2Δ-Sh3). (**I**) Dot plot showing the difference in cell proliferation in the 3T3 fibroblasts overexpressing WT-Sh3- vs ASD2Δ-Sh3/GFP-control with and without Fasudil (HF). Data is depicted as the BrdU incorporation by cells as percent of GFP-control. (**J**) Bar graph representing the relative mRNA expression of markers of fibrosis and EMT in 3T3 cells (WT-Sh3- vs ASD2Δ-Sh3). [Line and whiskers indicate mean ± SEM; Unpaired T-test *p < 0.05, **p < 0.01, ***p < 0.001, ****p < 0.0001].

To screen for the profibrotic domain in Shroom3(27), we first generated deletion mutants of Shroom3, each without one of the three known consensus domains (PDZ, ASD1, or ASD2) [1B]. The ASD2-domain is known to bind and activate Rho-associated coil-coiled containing protein kinases (ROCK1, and ROCK2)(27). We confirmed that ASD2-domain deletion (ASD2Δ-Sh3) abolished ROCK binding in HEK 293 tubular cell lines vs other deletional mutants [Fig 1C]. We then screened each domain-deletion variant, against WT-Sh3 (whole Shroom3) using a TGFβ1/Smad reporter assay in HEK-293 blue^TM^ cells. In this screening assay which detects secreted alkaline phosphatase in response to Smad3/4 signaling, ASD2Δ-Sh3 inhibited TGFβ1 signaling reporter at baseline and in the presence of TGFβ1[S1B-C]. In HEK-293 cells, we co-transfected SMAD3/4-reported luciferase constructs with either WT- or ASD2Δ-Sh3 plasmids and analyzed TGFβ1-signaling with/without TGFβ1. Again, ASD2Δ-Sh3 construct significantly reduced TGFβ1-signaling vs WT-Sh3 [Fig S1C]. We also evaluated other tubular- (mIMCD) and fibroblast- (NIH3T3) -cell lines stably overexpressing either WT-Sh3- vs ASD2Δ-Sh3 and studied Rock-activation and pro-fibrotic signaling. ASD2Δ-Sh3 overexpression reduced phospho-MYPT (P-MYPT) & total ROCK1 vs WT-Sh3 and/or control vector [1D-E], altered the actin cytoskeleton [S1D-E] and reduced cell migration of mIMCD cells [1F-G] vs WT-Sh3. In mIMCD, transcripts downstream of Wnt/Ctnnb1- signaling, TGFβ1-related profibrotic markers and pro-inflammatory chemokines were all significantly reduced in ASD2ΔSh3, while WT-Sh3 demonstrated increases of these transcripts vs. control-vector cells [1H]. P-Mypt levels and gene-expression changes demonstrated that ASD2ΔSh3-cells slightly inhibited Rock-activation even when compared to control vector, suggesting a dominant negative effect of ASD2Δ-Sh3 variant on endogenous Shroom3. WT-Sh3 induced P-MYPT excess and transcript changes were reduced by Rock-inhibitor HF, suggesting these effects were mediated by Rock-activation [Fig S1F-G]. Next, in 3T3 fibroblasts overexpressing ASD2Δ-Sh3 or WT-Sh3 [Fig S1H], ASD2Δ-Sh3 also inhibited P-MYPT [Fig S1H], profibrotic marker production [1I], and proliferation [1J] vs WT-Sh3-overexpression. Hence, we observed a key role of ASD2 domain of Shroom3 on Rock binding/activation impacting cytoskeleton, cell migration, and profibrotic signaling in tubular-/fibroblast cells.

### 2. Generation and phenotyping of inducible, global, ASD2Δ-Sh3 overexpression mice

To evaluate the role of the ASD2-domain of Shroom3 *in vivo*, we generated novel Dox-inducible, transgenic mice with Flag-tagged, ASD2Δ-Sh3, that could be mated with global (CAGS-rtTA) or tissue specific-rtTA and compared these to intact Shroom3-overexpressing mice (WT-Sh3 mice) [shown in 2A].

We induced overexpression of Shroom3 variants in these lines by DOX feeding [See methods]. Among multiple transgenic lines initially generated, a founder line each for ASD2ΔSh3 and WT-Sh3 was selected for inducible- and optimal- (2-3 fold) overexpression of Flag-tagged Shroom3 variants in kidney cortex by 3 weeks (wks) DOX [Fig S2A-S2B]. By immunofluorescence(IF) [2B], in CAGS-rtTA mice, overexpression in glomerular, tubular and interstitial kidney cells was observed for each of these lines vs non-DOX fed littermates, with apical localization of Shroom3 in tubular cells (near lotus lectin positivity), as described before(28). By qPCR for exon specific primers for the human ASD2-domain, only WT-Sh3 mice showed overexpression of this motif as expected [Fig S2C]. These mice were born at normal rates, and inducible overexpression of ASD2Δ-Sh3 and WT-Sh3 by itself was not associated with weight loss up to 4-wks[2C]. WT-Sh3 mice tended to have higher BUN and Creatinine levels by 3-wks DOX [2D, S2D], but neither line showed increased albuminuria vs non-transgenic littermates[2E]. We isolated and cultured primary renal cells (PCRCs) of ASD2ΔSh3 and WT-Sh3 after DOX-induction(29), and observed significantly reduced migration of ASD2Δ-Sh3 PCRCs vs WT-Sh3 PCRCs [2F-G], as seen *in vitro* with stably overexpressing cell lines.

Parallelly, we developed Fyn-binding domain mutant-Shroom3 overexpressing, transgenic mice (FBD-Sh3; with -AxxA- in place of -PxxP- at positions 442-445 between the PDZ- and ASD1-domains [see Fig 1B](14)) [2A], which were also mated with CAGS-rtTA animals. We confirmed overexpression of flag-tagged Shroom3 mutant in kidney cortex upon DOX [2H], with overexpression in glomerular cells on IF[S2E]. While Dox feeding of adult CAGS-rtTA/ ASD2ΔSh3 or WT-Sh3 mice did not induce albuminuria [2E] or podocyte FPE, inducible overexpression of FBD-Sh3 induced albuminuria [S2F] & FPE[2I-J], vs WT-Sh3- and/or non-Dox-fed FBD-Sh3 -animals. Together, these data showed a *role for FBD-Sh3 in anti-proteinuric function of Shroom3 in podocytes, along with the lack of proteinuria or podocyte injury in global* ASD2ΔSh3 overexpression mice.

### 3. Dox-inducible, global ASD2Δ-Sh3 overexpressing mice show reduced TIF vs WT-Sh3 overexpressing mice in AAN model

To study the role of ASD2-domain of Shroom3 in TIF development, we induced <u>Aristolochic acid nephropathy model (AAN)</u> in ASD2Δ-Sh3 and WT-Sh3. AAN is a model of tubular injury followed by TIF. AAN was induced after transgene induction using a multi dose regimen [n=5 mice each; see methods; Fig 3A]. Mice were evaluated at 9-wks allowing for post-injury remodeling for TIF. At 9-wks, CAGS-rtTA/ASD2Δ-Sh3 showed reduced azotemia vs CAGS-rtTA/WT-Sh3 mice [3B, S3C]. Both groups showed increased albuminuria after AAN induction vs baseline, but without significant differences between groups[S3D]. CAGS-rtTA/WT-Sh3 mice correspondingly showed significantly increased TIF by Masson-trichrome stain [3C], increased Collagen-III [3D], and Collagen-I staining [3E] by IF [quantified in 3F-H, respectively]), vs CAGS-rtTA/ASD2Δ-Sh3. In both mouse lines, kidney expression of Shroom3 was confirmed by IF[S3A] and by qPCR (including ASD2-domain specific *SHROOM3* and mouse *Shroom3* primers) [S3B]. Interestingly, WT-Sh3 mice showed reduced LTL-positive tubules with (a) more pronounced Shroom3 staining in tubular cell bases (instead of LTL-adjacent pattern seen in uninjured animals), and (b) greater interstitial Shroom3 expression, while ASD2Δ-Sh3 showed better preserved LTL-positivity and apical Shroom3 staining in tubules, consistent with reduced tubular injury in ASDD-Sh3 at 9-wks [S3A]. Consistent with invitro data, lysates from injured and uninjured kidneys in ASD2Δ-Sh3 mice showed reduced p-MYPT and ROCK1 vs WT-Sh3 lysates [Fig S3E]

**Figure 2:**
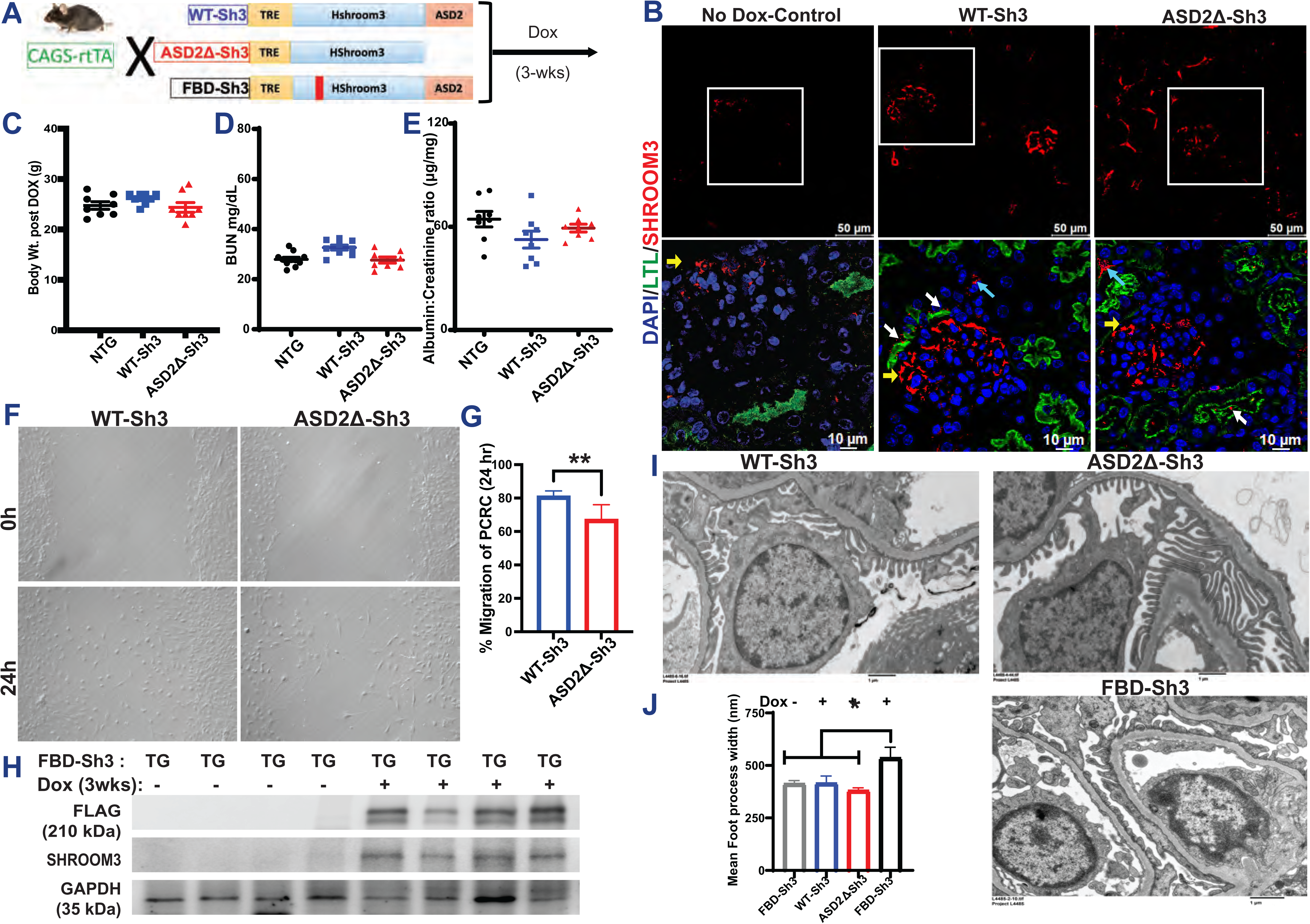
(**A**) Schema of generation of the Tetracycline-inducible global overexpression mice for WT-Sh3, ASD2Δ-Sh3 and FBD-Sh3. (**B**) Validation of the overexpression histologically by immunostaining for SHROOM3 (Red) in upper panel; Scale bar: 50 mM. The insets (lower panel) show co-staining with Lotus tetragonolobus lectin [LTL], Green and the white, yellow and blue arrows marking the tissue localization; Scale bar: 10 mM. Dot plots of baseline measurements of (**C**) Body weights (**D**) blood urea nitrogen (BUN) and (**E**) Albumin to creatinine ratio post Doxycycline induced overexpression in Non-transgenic (NTG)/WT-Sh3/ ASD2Δ-Sh3 mice (**F**) Phase contrast micrographs of wound closure in Primary culture renal cells (PCRCs) isolated from WT-Sh3/ ASD2Δ-Sh3 transgenic (TG) mice and the (**G**) Bar graph representing the percent migrated area (12 hpf lengths/well & 2 wells/line). (**H**) Representative immunoblots of whole kidney lysates from FBD-Sh3 mice probed for SHROOM3 and FLAG confirming overexpression on DOX induction. (I) Representative electron micrographs of glomeruli of WT-Sh3/ ASD2Δ-Sh3 vs FBD-Sh3 TG mice revealing the foot processes effacement in FBD-Sh3. (**J**) Bar graph depicting the mean foot process width [Line and whiskers indicate mean ± SEM; Unpaired T-test *p < 0.05, **p < 0.01].

**Figure 3:**
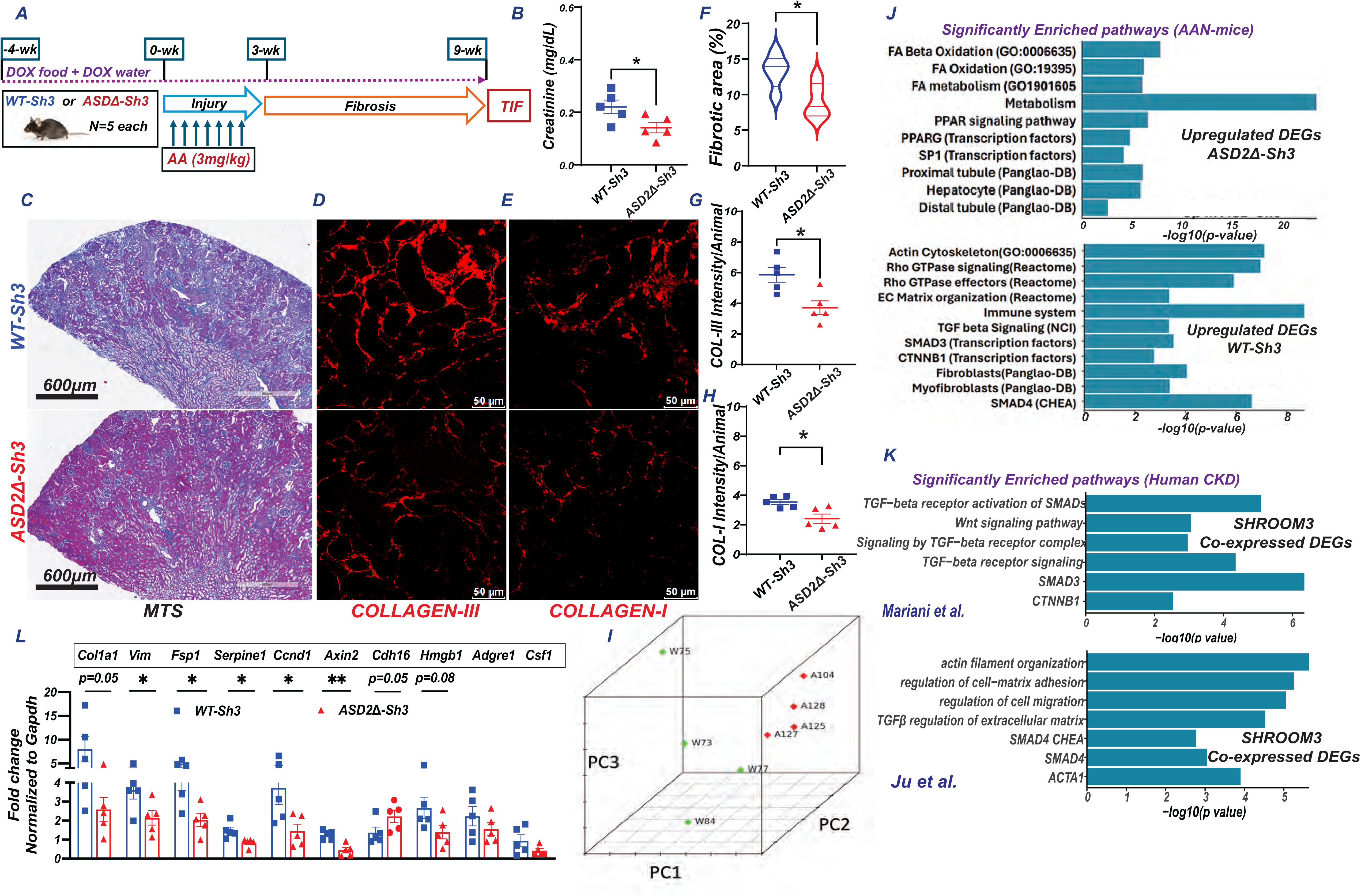
(**A**) Schema of AAN injury/TIF in CAGS-rTTA/WT-Sh3 or -ASD2Δ-Sh3 [n=5 each] (**B**) Dot plot of serum creatinine levels in AAN mice at 9-wks. Representative images of (**C)** slide-wide Masson’s Trichrome staining (MTS-4x) and (**D**) Collagen III and (**E**) Collagen I immunofluorescence to evaluate fibrosis in AAI-injured kidney. Scale bar: MTS- 600 μm; IF: 50 μm. Dot plots denoting the quantification of (**F**) fibrotic area (10 hpf per animal) and mean fluorescence intensities (12 hpf per animal) of (**G**) Collagen III and (**H**) Collagen I. (**I-K**) Bulk transcriptome evaluation by RNAseq was performed at 9-wks CAGS-rtTA- WT-Sh3 or -ASD2Δ-Sh3 mice (n=4 each). **(I)** Principal component analyses plot shows clustering between conditions Bar graphs show significantly selected enriched pathways from these analyses in **(J)** DEGs upregulated in ASD2Δ-Sh3 (upper panel) or WT-Sh3 (lower), while in **(K)** bar graphs show analogous pathways obtained from genes co-expressed with *SHROOM3* in 2 human CKD datasets. **(L)** Bar graph representing the relative mRNA expression (qPCR) of fibrotic/injury, inflammatory and tubular markers between transgenic lines. [Line and whiskers indicate mean ± SEM; Unpaired T-test*p < 0.05, **p < 0.01; DEGs=Differentially expressed genes].

To evaluate differentially expressed gene (DEG) between the kidneys of CAGS-rtTA/ASD2Δ-Sh3 & -WT-Sh3 during TIF, RNAseq was performed on total RNA from CAGS-rtTA/ASD2Δ-Sh3 & -WT-Sh3 kidneys at 9 wks [n= 4 vs 4 respectively; see methods]. A principal component analyses plot showed clustering of ASD2Δ-Sh3 & -WT-Sh3 transcriptomes demonstrated transcriptome-wide differences in TIF kidneys between these genotypes[3I]. Differential gene expression was analyzed after aligning reads with both mouse- (mm39) as well as aligning human orthologs of transcripts with human (GRCh38) transcriptome to facilitate downstream enrichment analyses [see methods]. Significant DEGs identified by DESeq2 from both alignments are tabulated in Table S1 and S2 respectively. In GRCh38 alignment, we identified 534 DEGs with a P-value threshold P<0.01[Fig S3F; Table S2]. Significantly upregulated DEGs in WT-Sh3 transcriptome (downregulated in ASD2Δ-Sh3) demonstrated significant enrichment of signals related to small GTPase/Rho-kinase activation, along with enhanced TGFβ1/Wnt-b-Catenin signaling, ECM production and fibrosis [3J]. Consistent signals of Rho-kinase activation, and fibrosis were also identified when the top 500 upregulated DEGS in WT-Sh3 mice by fold change were evaluated [P<0.05; Fig S3G]. Conversely, upregulated DEGs in ASD2Δ-Sh3 demonstrated signals of preserved proximal tubular cell homeostasis and metabolism, suggesting reduced injury [Fig 3J & S3F]. We also validated key mRNA signals by qPCR. Profibrotic gene transcripts, transcripts related to Wnt/Ctnnb1 signaling were significantly elevated in WT-Sh3 kidneys, while markers of tubular homeostasis were reduced [3L]. Proinflammatory transcripts (chemokines, immune cell related) also tended to be increased in WT-Sh3 vs ASD2Δ-Sh3 kidneys similar to data in WT-Sh3 mIMCD cell lines both by RNAseq and qPCR [Fig S3F &3L].

To compare these gene expression changes in WT-Sh3 overexpression with human CKD, we performed co-expression analyses to identify genes that significantly correlated with *Shroom3* from tubulo-interstitial transcriptomes of three human CKD cohorts within Nephroseq(R³0.4; 429±83 genes)(30–32). Enrichment analyses of co-expressed genes revealed significant enrichment of small-GTPase Rho-/ TGFβ1-/Wnt-beta Catenin- signaling consistent with data from WT-Sh3 overexpressing mice with TIF in our AAN model [Fig 3K/S3H vs Fig 3J/S3G] and showed clear translational relevance of our murine model.

### 4. Dox-inducible, global ASD2Δ-Sh3 overexpressing mice show reduced TIF vs WT-Sh3 overexpressing mice in UUO model

We evaluated a second model of TIF– unilateral ureteric obstruction (UUO) as shown in Fig 4A (n=4 vs 5 mice CAGS-rtTA/ WT-Sh3 and ASD2Δ-Sh3, respectively). UUO was induced after transgene induction with DOX as described before (see methods) (33), and obstructed kidneys were evaluated for TIF at 7 days. qPCR confirmed increases in human ASD2-domain specific primers in WT-Sh3 UUO kidneys [4B], while mouse Shroom3-specific primers remained similar between UUO kidneys of both transgenic animals [Fig S4A]. In the UUO model again, WT-Sh3 mice showed increased TIF by Trichrome staining[4C-D], and by Picrosirius red staining for Collagens [4E-F; see methods] vs ASD2Δ-Sh3 mice. Immunoblotting of cortical lysates confirmed increased phosphorylation of Smad3 in WT-Sh3 animals[4G-H]. Analogously, profibrotic, EMT-related, pro-inflammatory signaling gene transcripts were significantly increased in WT-Sh3 UUO kidneys vs ASD2Δ-Sh3 UUO kidneys [Fig 4I]. These data demonstrated mitigated TIF in a second model by global over-expression of ASD2-domain deleted Shroom3 compared to conditions of WT-Sh3 excess.

**Figure 4:**
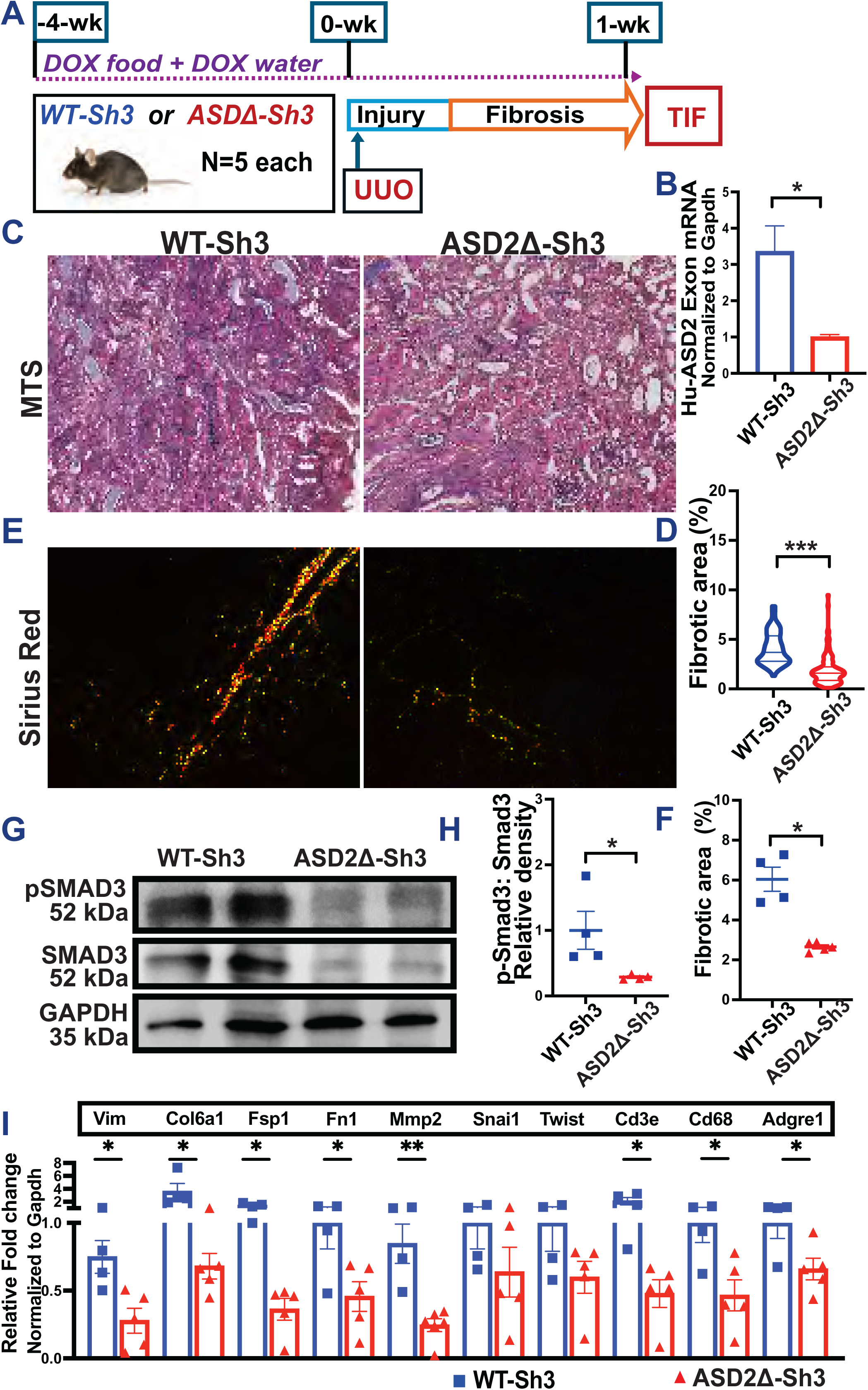
**(A)**TIF was induced by UUO model in CAGS-rtTA/WT-Sh3 or -ASD2Δ-Sh3 [n=4 vs 5]. **(B)** Bar graph representing the relative mRNA expression using ASD2-domain specific primers. **(C)** Representative images of (**C)** Masson’s Trichrome staining (MTS-20X) and (**E**) Picrosirius red (plane polarized light 20X) to evaluate TIF in UUO kidneys. **(D, F)** Dot plots quantify MTS and Sirius red among all mice, respectively (>10 hpf per animal). **(G)** Representative immunoblots of whole kidney lysates from UUO kidneys of WT-Sh3 or -ASD2Δ-Sh3 mice probed for P-SMAD3, SMAD3, GAPDH, and (H) dot plots show respective quantification of P-SMAD3:SMAD3 ratio by densitometry. **(I)** Bar graph representing the relative mRNA expression (qPCR) of fibrotic/injury, inflammatory and tubular markers between UUO kidneys between transgenic lines. [Line and whiskers indicate mean ± SEM; Unpaired T-Test *p < 0.05, **p < 0.01; ***p < 0.001; DEGs=Differentially expressed genes].

### 5. Dox-inducible, tubular-specific ASD2Δ-Sh3 overexpressing mice show reduced TIF vs WT-Sh3 overexpressing mice in AAN model

Based on the expression of Shroom3-Rock in tubular cells [Fig 1A], and the reduced TIF in post-tubular injury models, we crossed our Shroom3 transgenic mice with Pax8-rtTA lines for tubular specific over-expression. The mRNA expression analysis of kidney cortex of these Pax8-rtTA-Shroom3 mice confirmed overexpression of ASD2-domain specific sequences in WT-Sh3 mice vs other lines [Fig 5A] after DOX-induction. Dox-feeding did not induce increased creatinine and tended to slightly increase albuminuria and BUN in Pax8-rtTA/WT-Sh3 mice vs - ASD2Δ-Sh3 [Fig5B-C, S5A; n=8 each].

**Figure 5:**
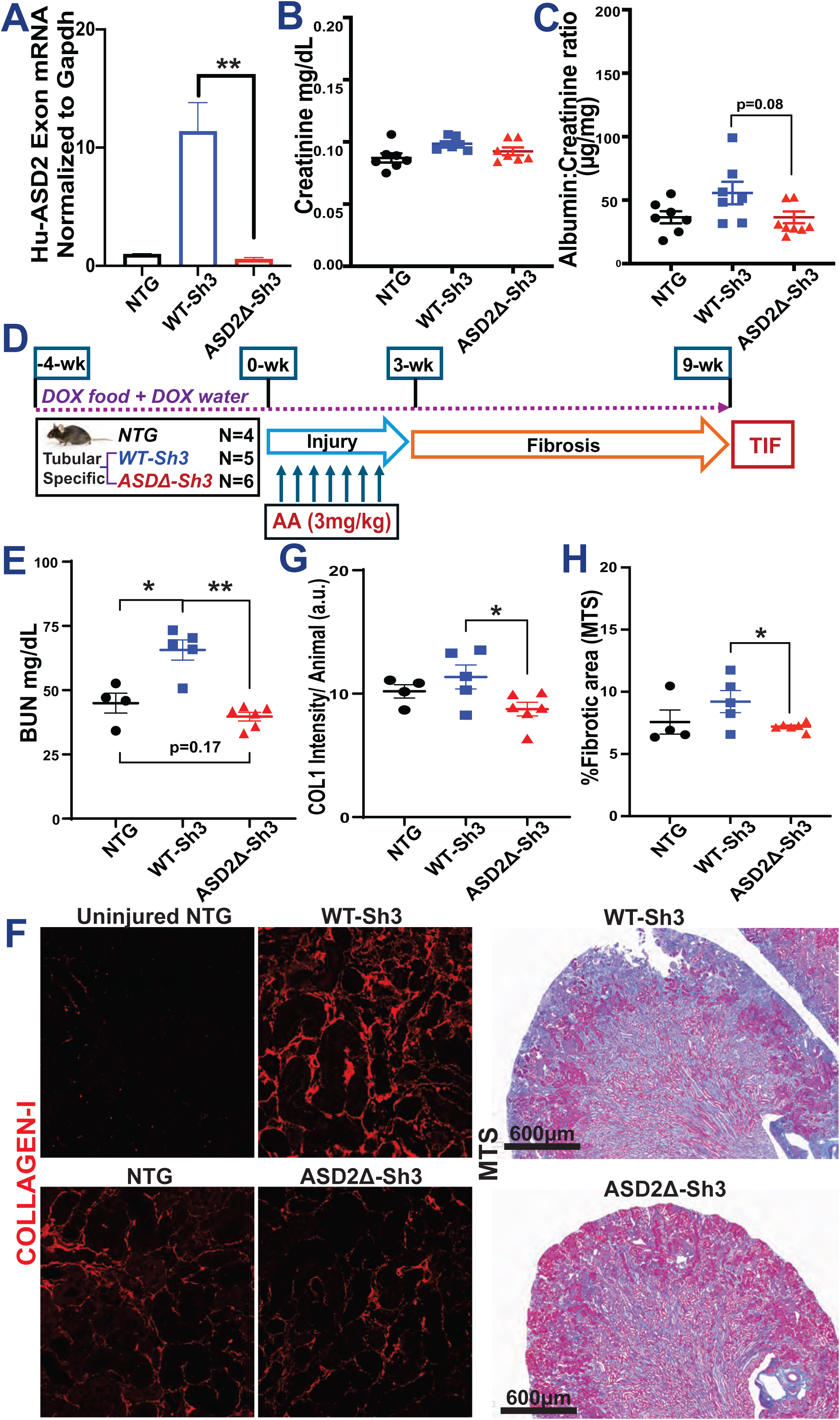
(**A**) Bar graph representing the relative mRNA expression by qPCR using ASD2-domain specific primers in tubular-specific (Pax8-rtTA) WT-Sh3 overexpression mice vs. Pax8-rtTA-ASD2Δ-Sh3 mice and non-transgenic controls. Dot plots of baseline levels of (**B**) serum creatinine and (**C**) urine albumin to creatinine ratios in DOX-treated Pax8-rtTA mice and non-transgenic mice. (**D**) Schema of generation of Aristolochic acid nephropathy (AAN) in tubular-specific overexpression WT-Sh3/ASD2Δ-Sh3 mice entailing injury followed by recovery phase with fibrosis. (**E**) Dot plot showing the BUN levels in AAI-injured Pax8-Shroom3 mutant mice at 9 wk. (**F**) Representative fluoromicrographs (20X) of Collagen I immunostaining and slide-wide photomicrographs of Masson’s Trichrome staining (MTS) to evaluate fibrosis in AAI-injured kidney. Dot plots denoting the (**G**) mean fluorescence intensities (12 hpf per animal) of Collagen I immunostained sections and (**H**) quantification of blue-stained area (10 hpf per animal) in Trichrome-stained sections. [Line and whiskers indicate mean ± SEM; Unpaired T-test*p < 0.05, **p < 0.01]

We then induced AAN after transgene induction (4-wks) to compare Pax8-rtTA/WT-Sh3 mice vs - ASD2Δ-Sh3 as well as Dox-fed non-transgenic littermates [n=6 vs 5 vs 4 mice respectively; see methods; Fig 5D]. At 9-wks, WT-Sh3 had greater azotemia [5E and S5C] than ASD2Δ-Sh3 or NTG animals. Immunofluorescence of AAN kidneys from Pax8-rtTA/WT-Sh3 mice & - ASD2Δ-Sh3 animals showed increased Shroom3 staining in tubular cells [S5B]. Similar to CAGS-rtTA animals, WT-Sh3 with greater injury showed more cytoplasmic staining while ASD2Δ-Sh3 tubules showed maintained apical localization with LTL-positive areas. WT-Sh3 mice showed increased Collagen-I staining by IF [5F and 5G; see methods], and TIF by Trichrome staining [5F and 5H] vs ASD2Δ-Sh3 mice/NTG. WT-Sh3 mice with greater overall injury from TIF also tended to have greater albuminuria [S5D].

Taken together these data confirm <u>a key role of Rock-binding ASD2-domain in the development of TIF and progressive kidney injury in a milieu of tubular cell excess of Shroom3 and show translational relevance of our findings to human CKD [Figure 6]</u>.

**Figure 6:**
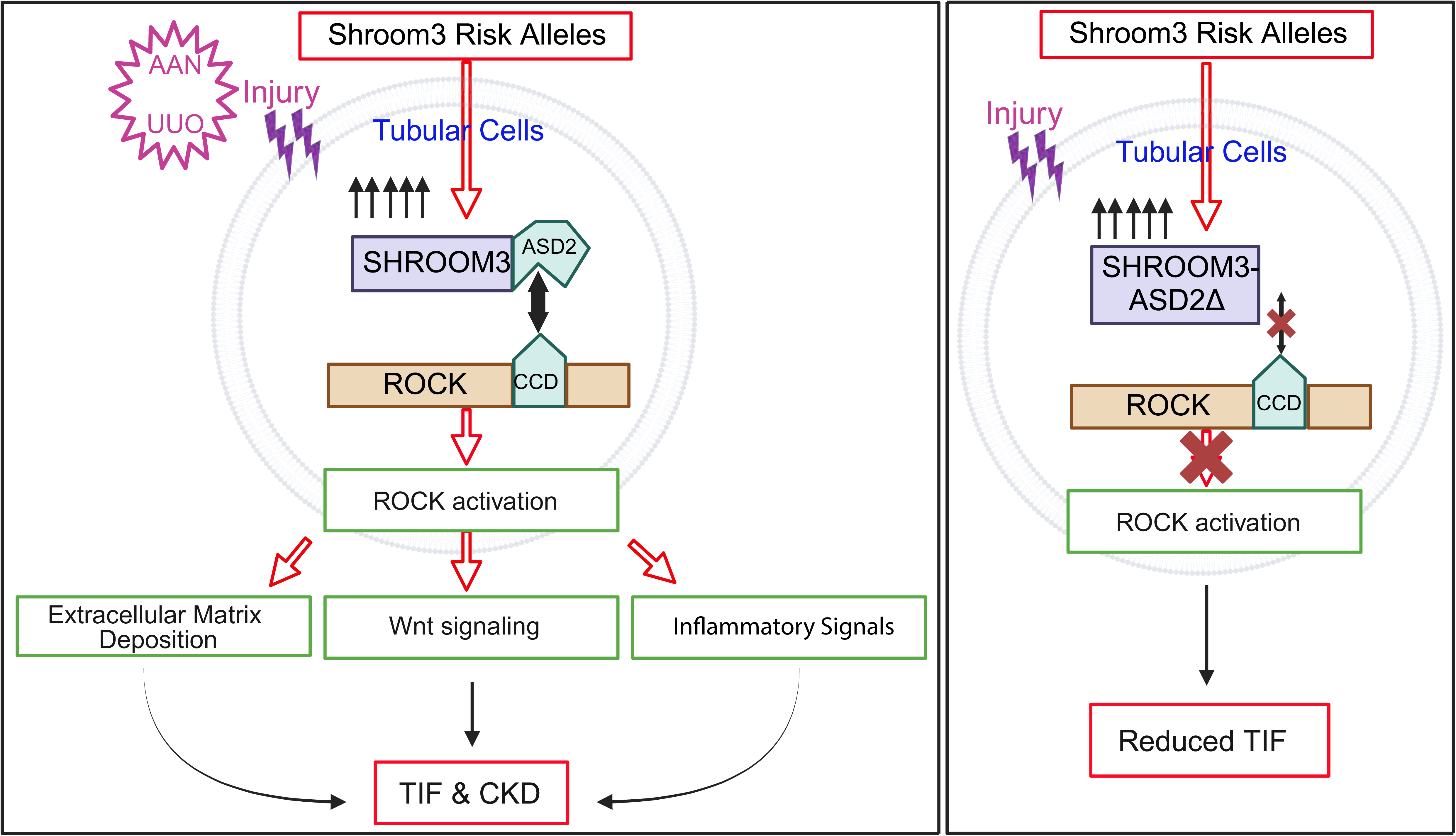
**Summary Schematic-** WT-Sh3 overexpressing mice mimic the Shroom3 excess in the kidneys of humans with the Shroom3 risk alleles which facilitates SHROOM3-ROCK1 and/or ROCK2 interaction in specific cell types *in vivo*. When kidney injury is induced (predominantly tubular injury) by either AAN or UUO the augmented ROCK1/2 activation within iPTs leads to increased TIF after injury attributable to increased Wnt/Ctnnb1 and TGFβ1 signals and downstream expression of pro-fibrotic markers, pro-inflammatory cytokines and chemokines (promoting fibro-inflammation). Together, these promote TIF. This is a mechanism that underlies faster progression to CKD in AKI patients with the Shroom3 risk alleles. Meanwhile, overexpressing ASD2Δ-Sh3 in the same cells *in vivo* abolishes ROCK1/2 interaction and reduces activation and reduced post-injury TIF in kidneys of ASD2Δ-Sh3 mice. Also, the data indicates a dominant negative effect of ASD2Δ-Sh3 overexpression over the endogenous Shroom3 activity. Our findings indicate the therapeutic potential of cell-specific inhibition of SHROOM3-ROCK1 and/or ROCK2 interaction for TIF and CKD progression in humans. (iPTs -injured proximal tubular cells) *Created with BioRender.com*

## Discussion

While multiple loci related to CKD have been identified by GWAS, few loci/genes from unbiased analyses have made significant progress towards precision medicine approaches. For instance, intronic variants in the Shroom3 gene have been repeatedly identified as associated with CKD (34, 35). Using detailed mechanistic studies, we previously showed that rs17319721 Shroom3 variant, when present in donor kidneys during transplantation, was associated with both increased intrarenal expression of *SHROOM3* and increased TIF (CADI-score) by 12-months(8). Independent work has also supported a regulatory role for rs17319721 in renal epithelial cells (9), and of other intronic Shroom3 variants as eQTLs specifically in proximal tubular cells(35). We also reported that shRNA-mediated Shroom3 knockdown *in vivo* mitigated TIF in a UUO model and identified crosstalk between TGFβ1- and Wnt/Ctnnb1 pathways mediated by Shroom3(8). These data supported the concept that targeting Shroom3 in patients or donor-kidneys with Shroom3 SNPs could be an exploitable therapeutic strategy. However, multiple reports have since showed that Shroom3 has potentially beneficial roles in podocytes both during development (12, 36), and in adult animals (14), and non-selectively antagonizing Shroom3 protein would incur the unacceptable cost of proteinuria. Our group evaluated the role of Shroom3 in podocyte function in adult animals and reported a previously unidentified Fyn-binding motif in Shroom3. We showed that adult mice with podocyte-specific Shroom3 knockdown phenocopied Fyn-KO mice, demonstrating that this domain played a key role in actin cytoskeletal organization and adult podocyte homeostasis(14).

In the current series of experiments, we tested the hypothesis that a distinct domain, i.e. the ASD2 (Rock-binding) domain in Shroom3 plays the key role in mediating the association of high-expressor Shroom3 SNPs with TIF identified in data evaluating CKD. Here, we first screened deletional variants without Shroom3 consensus domains *in vitro* using TGFβ reporter assays and identified mitigation of TGFβ1/Smad signaling in ASD2Δ-Sh3 variant. In tubular-cell lines ASD2Δ-Sh3 overexpression with reduced Rock/Rock activation also showed significantly reduced profibrotic and proinflammatory signaling vs WT-Sh3 overexpression. Based on ScRNAseq data showing that co-expression of Shroom3 and both Rock paralogues occurred in fibroblasts in addition to injured tubular cells, we also tested ASD2Δ-Sh3- vs WT-Sh3- overexpression in fibroblast cell lines and observed reduced profibrotic signaling in ASD2Δ-Sh3-fibroblasts – albeit with lower effect sizes. Based on these data showing that abolishing Shroom3-Rock interaction, Rock activation and consequent downstream Rock signaling, could mitigate profibrotic and proinflammatory signals that arise in the context of Shroom3 excess, we tested this idea *in vivo* using novel transgenic mice that were ASD2Δ-Sh3 vs WT-Sh3. Consistently in two models of TIF, we identify that global Shroom3 excess mediated increases in TIF & CKD are protected when ASD2Δ-Sh3 variant, i.e. with inability to bind and activate Rock1 or-2, is overexpressed in place of WT-Sh3. Our results are consistent with prior reported benefits of ROCK-inhibitors in experimental TIF (37). Transcriptomic data confirmed the activation of ROCK-, TGFβ1- & Wnt/Ctnnb1- mediated profibrotic signaling in the presence of CKD with excess WT-Sh3 (vs ASD2Δ-Sh3 variant), and comparison to human CKD transcriptome using co-expression analyses showed relevance of these findings to human TIF kidneys in CKD. Finally, using tubular-specific overexpression (Pax8-rtTA) mice, we isolated the injured tubular cells as the source where excess Shroom3-Rock interaction during CKD promoted injury, and where antagonizing Shroom3-Rock interaction in high expression states could reduce injury and TIF. Critically, while ASD2Δ-Sh3 and WT-Sh3 variants did not induce proteinuria or podocyte injury, FBD-Sh3 overexpression mice induced podocyte injury, showing distinct motif-specific effects from SHROOM3 occurring in a cell-type specific manner.

Our work has several implications for CKD. First, our experimental work ascribes a detailed mechanism to the often-identified association of highly prevalent intronic cis-eQTL SNPs in Shroom3 (Allele frequencies ∼ 0.29-0.4) with CKD in the general population. Next, this mechanism likely has application to both native and allograft CKD occurring in donors with risk alleles (8, 38). Critically, the protein-protein interaction-based, pro-injury signaling mechanism that we describe here may potentially allow for the development of pharmacologic tools that could inhibit Shroom3-Rock P2I and mitigate CKD progression(39).

Our data has some limitations. Our *in vitro* data also suggests a potential role for Shroom3-Rock interaction in fibroblast function. The specific contribution of Shroom3-Rock interaction in vivo in <u>fibroblasts</u> (vs the contribution of tubular cell overexpression) during TIF/CKD development needs to be evaluated in future work including fibroblast-specific over-expression of Shroom3 variants. From the benefit in azotemia and TIF we observed in Pax8-rtTA animals, the contribution of fibroblast Shroom3-Rock could be inferred to be lower, although this needs experimental testing. Interestingly, in contra-distinction to multiple reported <u>adverse associations of Shroom3-SNPs with CKD, a</u> potential <u>protective role for a linked intronic Shroom3-SNP in AKI</u> in humans was reported recently(40), and increased AKI also occurred in heterozygous global Shroom3-knockout mice(28). Hence, clearly delineating the role of increased Shroom3-Rock interaction during acute injury vs in post-injury TIF will be necessary before therapeutic use to avoid any potential harm during AKI. Although we did not conduct such time course experiments here, it must be noted that we continuously induced ASD2Δ-Sh3 throughout the injury phase and still observed benefit in TIF. Finally, using transgenic models, any Rock-independent effects of the Shroom3-ASD2 domain on TIF needs to be ruled out. Along these lines, while Rock1- & 2 show near ∼90% homology, these have non-redundant cell-specific and temporal expression in kidney cells during injury(41), and whether inhibiting one or both kinases is essential for anti-fibrotic effect is unclear. All these mechanistic studies are critical for translational potential and the ultimate development of Shroom3-Rock interaction inhibitors to treat CKD related to Shroom3-excess & Shroom3 SNPs.

In summary, our data reports a critical role for Shroom3-Rock interaction in injured tubular cells as a mechanism underlying increased TIF in situations of Shroom3 excess. The absence of the ASD2 domain reduces the ROCK activation and subsequent pro-fibrotic and pro-inflammatory signals and ECM deposition to ameliorate TIF progression. Our work unravels a plausible and targetable pathway that is specifically applicable to patients with progressive kidney disease who harbor the intronic enhancer Shroom3 SNPs that associate with CKD in GWAS and could have greater applicability.

## Methods

### Cell culture

The mIMCD-3 cell line was generously gifted by Dr. Stefan Somlo (Yale School of Medicine, New Haven, USA). The NIH3T3 and HEK293T cell lines (kindly gifted by Dr Lloyd Cantley, Yale School of Medicine, New Haven, USA). The former cells and the latter two cells were cultured in DMEM/F-12 and DMEM respectively, supplemented with 10% FBS and 1% penicillin-streptomycin and maintained at 37°C in a humidified atmosphere containing 5% CO2.

### Transient Overexpression studies

Human SHROOM3 mutant constructs (GenScript) with deletions of PDZ, ASD1 or ASD2 domains, as well as WT were transfected onto 293T cells using PolyJet transfection (SignaGen Labs) as previously described (8). All constructs used had a V5-tag.

### Luciferase assay

HEK293T cells were cotransfected with SBE4-Luc (0.5 μg) and pRL plasmids (0.2 μg) as 5 μg of either of the SHROOM3 mutant constructs, using the PolyJet Transfection Kit according to the manufacturer’s instructions (SignaGen Laboratories). The transfected cells were treated with 5 ng/ml TGFβ1 or vehicle for 15 min and Luciferase activities were measured using the Dual-Luciferase Reporter Assay Kit (E1910; Promega).

### Generation of stably-transfected cell lines

HEK293T cells were used as producer cells for lentiviruses. The cells were transfected with 5 µg each of the GFP, WT- or ASD2Δ-Shroom3 lentiviral plasmid (pLenti CMV/TO Puro DEST), pPACK, and pVSV using PolyJet transfection reagent (SignaGen Labs) to generate mammalian VSV-pseudotyped lentiviral expression constructs as previously described (8). The media containing lentiviral constructs were collected, filtered and concentrated using Lenti-X Concentrator (#631231;Takara) following the manufacturer’s protocol. These lentiviral constructs were used to infect mIMCD and 3T3 cells and the stably overexpressing cells were selected by puromycin (#ant-pr-1, InvivoGen) treatment.

### Western blot

Mouse kidney tissues, cell lines and primary renal cells were lysed using RIPA buffer added with Halt Protease and Phosphatase Inhibitor (#78440, Thermo Scientific,) as mentioned elsewhere (24300173). Briefly, the tissue or cells were homogenized in the lysis buffer using mechanical digestion (Precellys homogenizer, Bertin technologies) and incubated on a rotator at 4°C for 30 min. The lysate was centrifuged at 12000xg at 4°C for pelleting down the undigested cell debris. The protein concentrations of the supernatant lysates were determined by BCA assay (#23225, Pierce-Thermofisher). Loading buffer were then added and the mixtures were denatured under 95°C. The proteins were separated on 4-15% or 10% gels (Bio-Rad) by SDS-PAGE and transferred to an Immobilon-P membrane (Millipore,) followed by blocking with 5% BSA or 5 % skim milk. The primary antibodies used were: anti-MYPT(Cell Signaling), anti-phosphoMYPT-T696(), anti-ROCK1(Cell Signaling), anti-ROCK2 (#8236, Cell Signaling), anti-HSP-90 (#4877, Cell Signaling), anti-SHROOM3 (#SAB3500818, Millipore-Sigma; #LS-C679459, LSBio), anti-V5 (#V8012, MilliporeSigma), anti-FLAG (#A8592, Millipore-Sigma), anti-β-ACTIN (#A5441, MilliporeSigma), anti-AMPK(#2532, Cell Signaling), anti-phosphoAMPK-T172 (#2535, Cell Signaling). The secondary antibodies used for detection included horseradish peroxidase (HRP)-conjugated anti-rabbit and anti-mouse antibodies, with a dilution ranging from 1:8000 to 1:12000. Image Studio Lite Ver5.2 for western blot image acquisition. Densitometry was performed on images of Western blots using Image J software.

### Immunoprecipitation

PCDEST SHROOM3 (PDZΔ, ASD1Δ, ASD2Δ and WT) and GFP (control vector) overexpressing 293T cells were generated in 15 cm culture dishes. Protein lysates post 48 hours transfection and 15 min TGFβ (5ng/mL) treatment were immunoprecipitated with Anti-V5-tag mAb-Magnetic Beads (MBL #M167-11) and run on PAGE gels as described elsewhere (42).

### Immunocytochemistry-Immunofluorescence

WT-Sh3 and ASD2Δ-Sh3 overexpressing mIMCD3 cells were plated in 12-cm wells on cover slips, serum deprived for 8h and treated with Dmso or Fasudil (#S1573, Selleckchem) for 24 h. The treated cells were fixed and permeabilized (36% HCHO, 0.1% TritonX100 in PBS). The cells were blocked with 3% BSA/10% normal goat serum followed by incubation with primary antibody (1:100) against ROCK1 (#PA5-22262; Thermo Scientific). F-ACTIN staining for observing cytoskeletal changes in SHROOM3-transfected cells was done using 100nM Acti-stain 488 phalloidin (#PHDG1, Cytoskeleton Inc.) along with Anti rabbit Alexafluor 568 secondary antibody (1:300) (Invitrogen) in 1% BSA-PBS.

### Immunohistochemistry-Immunofluorescence

For immunostaining, 5 μm sections of formalin-fixed kidney tissues (processed at Yale Pathology Tissue Services (YPTS) facility were deparaffinized and subjected to antigen retrieval (Retrievagen (pH 6), BD Biosciences,) to unmask the antigens, followed by incubation with primary antibodies (Shroom-3 LS-Bio, #LS-C679459,1:100; Fluorescein LTL, Vector lab-#FL1321-2, 1:200; FLAG, Millipore-Sigma, #F1804,1:400; Collagen-I/III Southern Biotech, #1310-01/#1330-01, 1:100). Corresponding Goat anti-Rabbit Secondary Antibody, Alexa Fluor™ 488, #A-11008; Donkey anti-Goat Secondary Antibody, Alexa Fluor™ 594, #A32758 were used.

### Murine inducible global WT-Shroom3, ASD2Δ-Shroom3 and FBD-Shroom3 overexpression models

Plasmid constructs of WT, ASD2Δ and FBD Shroom3 (under TRE) tagged with FLAG were co-transfected with plasmid construct with reverse tetracycline transactivator (rtTA) under a CAGS promoter to generate inducible expression models. The resultant 3 fused plasmids were purified and used for transfecting mouse embryonic stem cells followed by generation of founder mice lines by the mouse core facility. For each construct, 5-7 founder lines were generated harboring random insertions in the genome. The mice from each founder line were tested for optimal (2-3 fold) and inducible overexpression in kidney- and tail- tissue post DOX induction. Founder mice of either gender (6-8 wks old) were provided DOX feed and DOX water for 4 wks and tested for mRNA by RT-qPCR and protein overexpression by Western blot. One founder line with the optimal overexpression for each of the Shroom3 mutants were used for *in vivo* experiments.

### Tubular specific WT-Shroom3 and ASD2Δ-Shroom3 overexpression models

The WT-Sh3 and ASD2Δ-Sh3 mice were crossed to mice expressing Pax8-rtTA for tubular-specific overexpression under DOX induction.

### Injury models

In the Aristolochic acid nephropathy (AAN) model, all mice received intraperitoneal injections (3mg/kg) of aristolochic acid (AAI, MilliporeSigma) every third day for 3 wks for a total of 8 injections and were euthanized 6 wks after the last injection (43, 44). In the Unilateral ureteral obstruction (UUO) model (45), mice were anesthetized with ketamine/xylazine and maintained on a 37°C heat pad during surgery. The right kidney was exposed via a flank incision, the ureter was ligated, and the mice were euthanized after 1 week. The kidney tissues were collected for histology, immunostaining for fibrosis markers, RNA isolation for RT-qPCR, and protein extraction for Westerns.

### Reverse transcription qPCR

Intron-spanning primer sets were designed for all assayed genes using Primer-BLAST (NCBI), and PCR amplicons were confirmed by both melting curve analysis and agarose gel electrophoresis. *Gene* expression was assayed by RT-PCR (Applied Biosystems 7500). Amplification curves were analyzed using the automated 7500 software platform via the ΔΔCt method. *Gapdh* was used as endogenous control. See Table S3 for primer sequences.

### Electron Microscopic analysis

Glutaraldehyde fixed samples were processed as previously described (46). The electron micrographs were captured at Electron Microscopy & Cryo Electron Microscopy facility in Center for Cellular and Molecular Imaging (CCMI), Yale. Average foot process width was determined as described elsewhere (47)

### Cell Proliferation Assays

Cell lines overexpressing the Shroom3 variants were plated on to 96 well plates and BrdU incorporation was performed according to the manufacturer’s protocol (#QIA58, MilliporeSigma).

### Primary Cultured Renal Cells

The primary renal cells were isolated from mouse kidneys and cultured for 5 days (26). Briefly, kidneys were harvested, minced, and digested in Liberase enzyme (0.5 mg/mL) with DNase I (100 mg/mL) (#5401151001, #10104159001, Roche Diagnostics) and 0.5mM MgCl₂ for 30 minutes. The resulting cell suspension was filtered, centrifuged, and incubated with RBC lysis buffer (#Invitrogen) for 3 min followed by washing with HBSS. The resultant cell pellet was resuspended in DMEM with 10% FBS and 1% penicillin-streptomycin and seeded into culture dishes. After 5 days of expansion and media changes, primary renal cell cultures (P0 PCRCs) which are 75% proximal tubules, were harvested and cryopreserved.

### Scratch Wound Assay

PCRCs or mIMCD3 cells expressing Shroom3 constructs were seeded in a 12-well culture plates at a density of approximately 70-80% confluence and incubated overnight at 37°C in a 5% CO₂ atmosphere. A sterile 10μl pipette tip was used to create perpendicular-line scratches across the cell layer, ensuring uniform width and depth. The plates were gently washed with sterile PBS to remove non-adherent cells. Fresh culture medium without FBS was added to each well, with or without experimental treatments. Images of the scratch area were captured at time zero (T0) using a microscope. Plates were then incubated as before. At 24h images were taken of the scratch area to observe cell migration. The scratch area was analyzed using ImageJ software (NIH, Bethesda, MD) to determine the percentage of wound closure over time.

### Bulk RNA-seq analyses

DNase treated RNA extracted from AAN-kidneys of WT-Sh3 and ASD2Δ-Sh3 (n=4) were depleted of rRNA and underwent quality control (QC) analysis at the Yale Center for Genome Analysis (YCGA) followed by library prep and Poly(A) RNA sequencing (NovaSeq, Illumina). The raw Illumina RNAseq paired-end data were quality controlled and adapter-trimmed using FastQC version 0.12.1 (48) and BBDuk version 39.11 (49, 50). We mapped the RNAseq data to reference genome using STAR version 2.7.11a (51). For read alignment, we used both the mouse reference genome (GRCm39) and the human reference (GRCh38). By mapping to human reference, we obtained the gene expression profiles in human orthologs. We computed the read counts in gene sets from mapped data using the Subread R package, FeatureCounts (52). To calculate differential gene expressions, we used DESeq2 version 1.44.0 (53). The principal component analysis (PCA) was computed using the DESeq2 plotPCA function.

### Statistical Analysis

Statistical analyses were performed using GraphPad Prism 10 (Dotmatics, CA). Data are presented as the mean ± SEM. Unpaired t-tests or Mann–Whitney tests were performed for univariate comparisons between two groups. One-way ANOVA (with Tukeys post hoc test) was used while comparing more than 2 groups. Statistical significance was considered with two-tailed P<0.05.

## Acknowledgements

MCM acknowledges funding from the NIH (grants R01DK122164, R01DK132274, and R21AI178705), and the Department of Defense, grants HT94252310454 and HT94252310441. MCM also acknowledges research support from the Blavatnik fund at Yale (Accelerator award) and a pilot award from CTSA Grant UL1 TR001863

## Supplementary Figure Legends

**Figure S1** (**A**) Dot plot representing *Shroom3, Rock1, Rock2* expression in multiple types identified in Day7 and Day30 IRI kidney (Unilateral Ischemia Reperfusion) vs. Sham Control kidney (Leyuan Xu and Lloyd Cantley). (**B**) Representative immunoblots of SHROOM3 and βACTIN using lysates from TGFβ- and vehicle-treated HEK293T cells overexpressing different Shroom3-variant constructs (top). Dot plots denoting the relative Smad reporter response of these cells in the presence and absence of TGFβ measured as the level of Secreted embryonic alkaline phosphatase (SEAP) produced in the culture supernatant (bottom). (**C**) Bar graph representing the Luciferase assay confirming the pronounced response to TGFβ by WT-Sh3 overexpressing HEK293T cells and the absence of this response in ASD2Δ-Sh3 cells. (**D**) Representative fluoromicrographs of immunostaining of ROCK1 and F-actin in WT-Sh3/ASD2Δ-Sh3 overexpressing mIMCD cells in the presence/absence of Fasudil (HF). (**E**) Violin plots denoting the quantification of the mean fluorescence intensities of F-actin and ROCK immunostaining, normalized to the total nuclei count in 5-8 hpfs per duplicate (n=2). Bar graphs depicting (**F**) the densitometry measurements for pMYPT1: βACTIN in WT-Sh3-mIMCD with/without HF treatments and (**G**) relative gene expression of a fibrosis marker and a cytokine which gets reduced by HF treatment in WT-Sh3-mIMCD. (**H**) Representative immunoblots of V5-tag (SHROOM3), pMYPT1, ROCK1 and HSP90 in 3T3 fibroblasts (WT-Sh3 vs. ASD2Δ-Sh3). *Line and whiskers indicate mean ± SEM; *p < 0.05, **p < 0.01, ***p < 0.001*

**Figure S2** (**A**) Immunoblot of FLAG tag to confirm overexpression of SHROOM3 *in vivo.* (**B**) Bar graphs denoting relative mRNA expressions of *Shroom3* in TG mice post DOX-induction. (**C**) Dot plot of mRNA levels of *Shroom3* exon harboring the ASD2-domain in WT-Sh3 mice vs. ASD2Δ-Sh3, post DOX-induction. (**D**) Dot plot of baseline serum creatinine levels in the DOX-treated mice. (**E**) Representative fluoromicrographs of SHROOM3/LTL immunostaining confirming overexpression in FBD-Sh3 mice post DOX-induction. (**F**) Bar graph representing urine albumin to creatinine ratios in FBD-Sh3 mice compared to other Shroom3-TG mice. *Line and whiskers indicate mean ± SEM; *p < 0.05*

**Figure S3** (**A**) Representative fluoromicrographs (40X and inlay) of SHROOM3/LTL immunostaining in AAN-kidney sections of global overexpression mice showing the cellular localization post-injury. (**B**) Bar graph denoting relative mRNA expressions of *Shroom3* exon harboring the ASD2-domain in AAI-injured kidneys of WT-Sh3 vs. ASD2Δ-Sh3. Dot plots depicting (**C**) the BUN levels and (**D**) Urine albumin to creatinine ratios in the AAN mice. (**E**) Immunoblots of pMYPT1, MYPT1, ROCK1 and βACTIN with AAI-injured and non-injured kidney lysates. (**F**) Volcano plot representing significant DEGs identified by DESeq2 from GRCh38 alignment for RNA sequencing dataset from AAN mice. Bar graphs representing the significantly enriched pathways identified by EnrichR analysis of (**G**) the top 500 upregulated DEGS in WT-Sh3 AAN-kidneys by fold change (p<0.05) and (**H**) co-expression analyses of enriched genes from another tubulo-interstitial transcriptome of human CKD cohorts within Nephroseq, which correlated with *Shroom3* expression. *Line and whiskers indicate mean ± SEM; *p < 0.05*

**Figure S4** (**A**) Dot plot denoting relative mRNA expressions of mouse endogenous *Shroom3* in UUO kidneys of WT-Sh3 vs. ASD2Δ-Sh3 mice. *Line and whiskers indicate mean ± SEM*

**Figure S5** (**A**) Dot plot showing the BUN levels in DOX-treated Pax8-rtTA mice and non-transgenic mice. (**B**) Representative fluoromicrographs of SHROOM3/LTL immunostaining in AAN kidneys of Pax8-rtTA- WT-Sh3/ASD2Δ-Sh3. Dot plots of (**C**) serum creatinine levels (**D**) urine albumin to creatinine ratios in the AAN mice. *Line and whiskers indicate mean ± SEM; *p < 0.05*

## Notes

### Competing Interest Statement

The authors have declared no competing interest.

