## Supplementary figures and images for "Shroom3-Rock interaction and profibrotic function: Resolving mechanism of an intronic CKD risk allele"

### Supplemental Figures

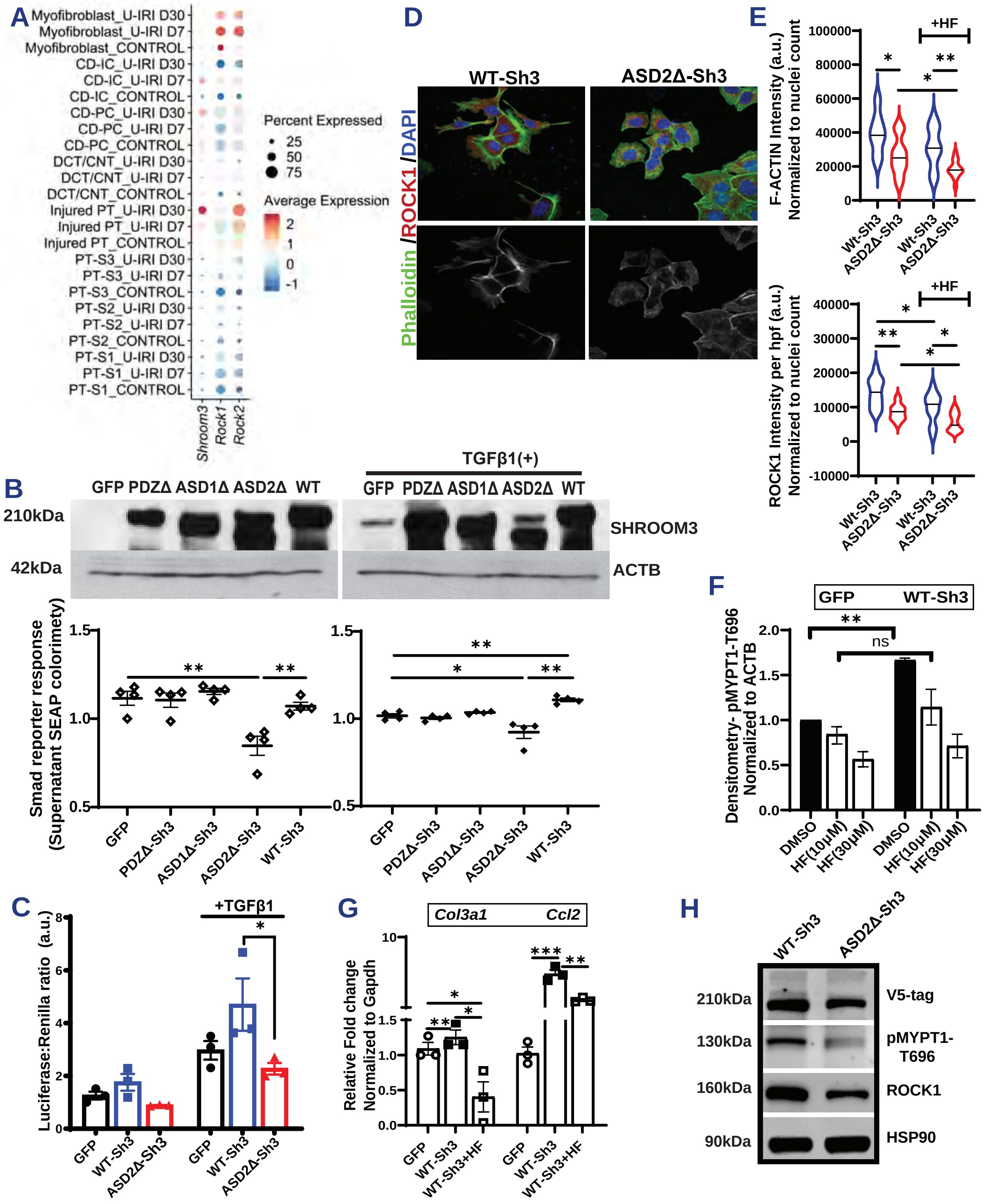

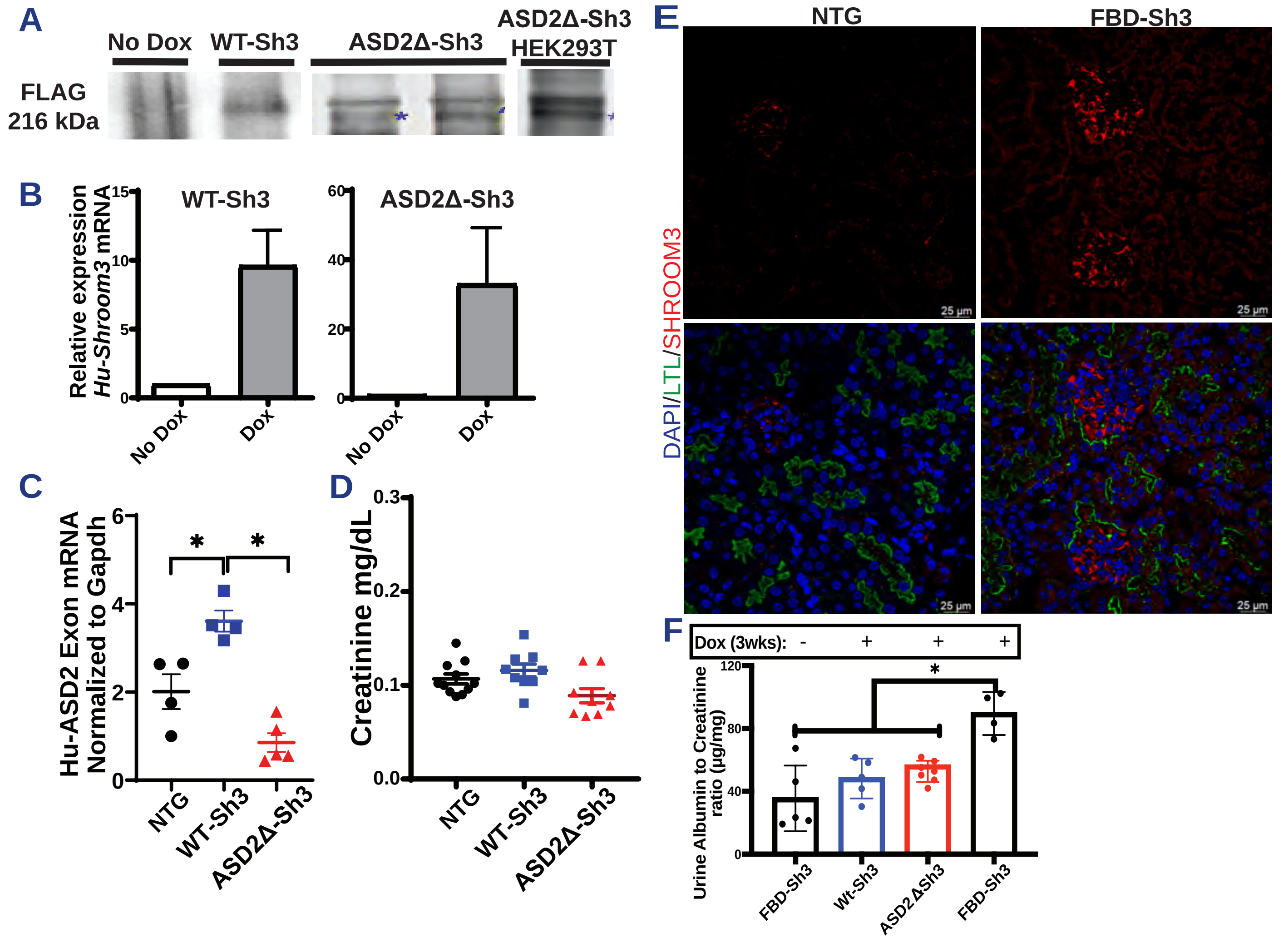

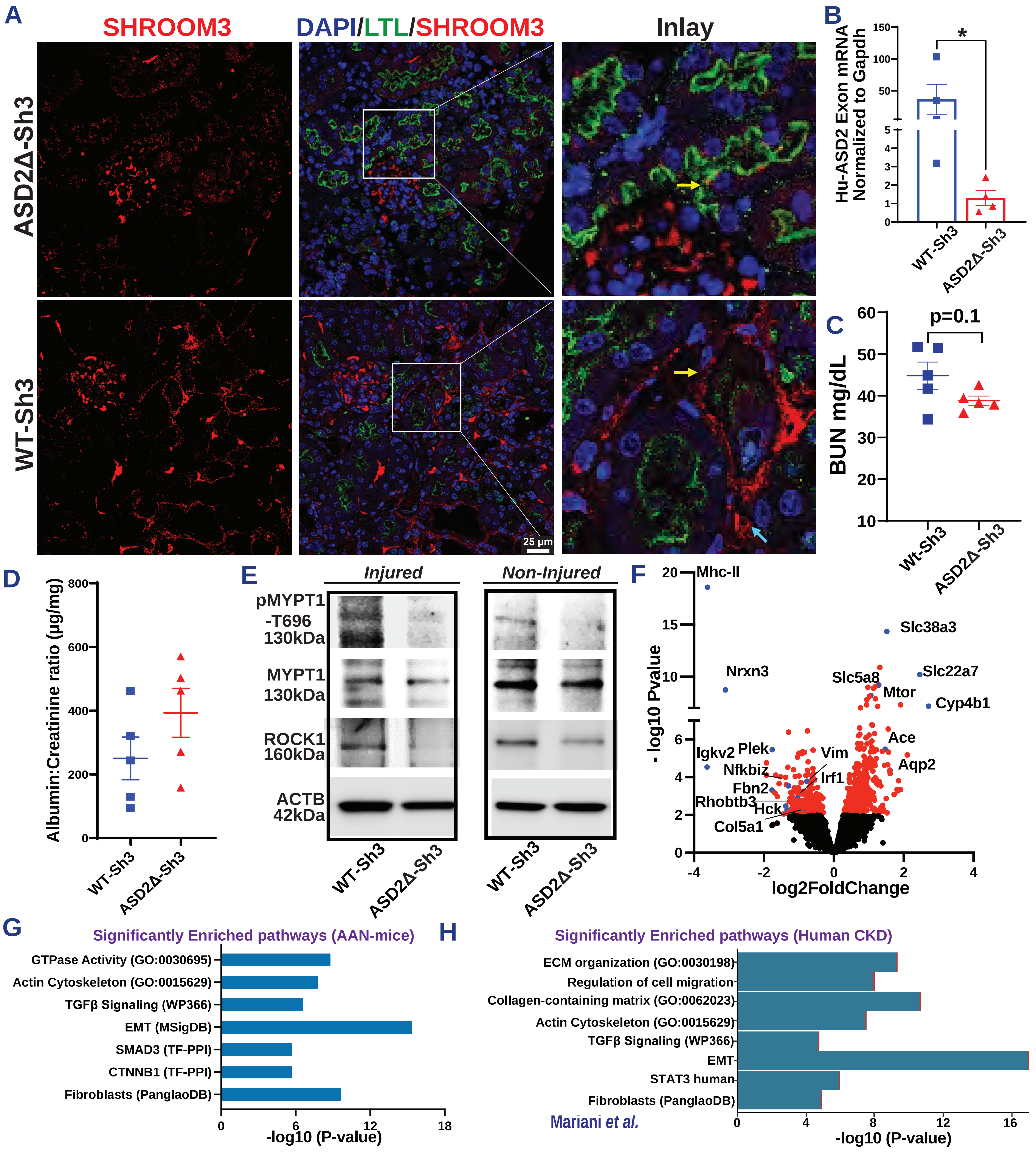

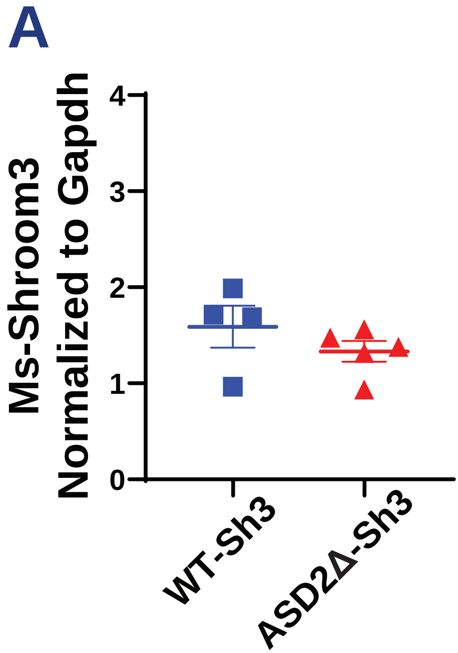

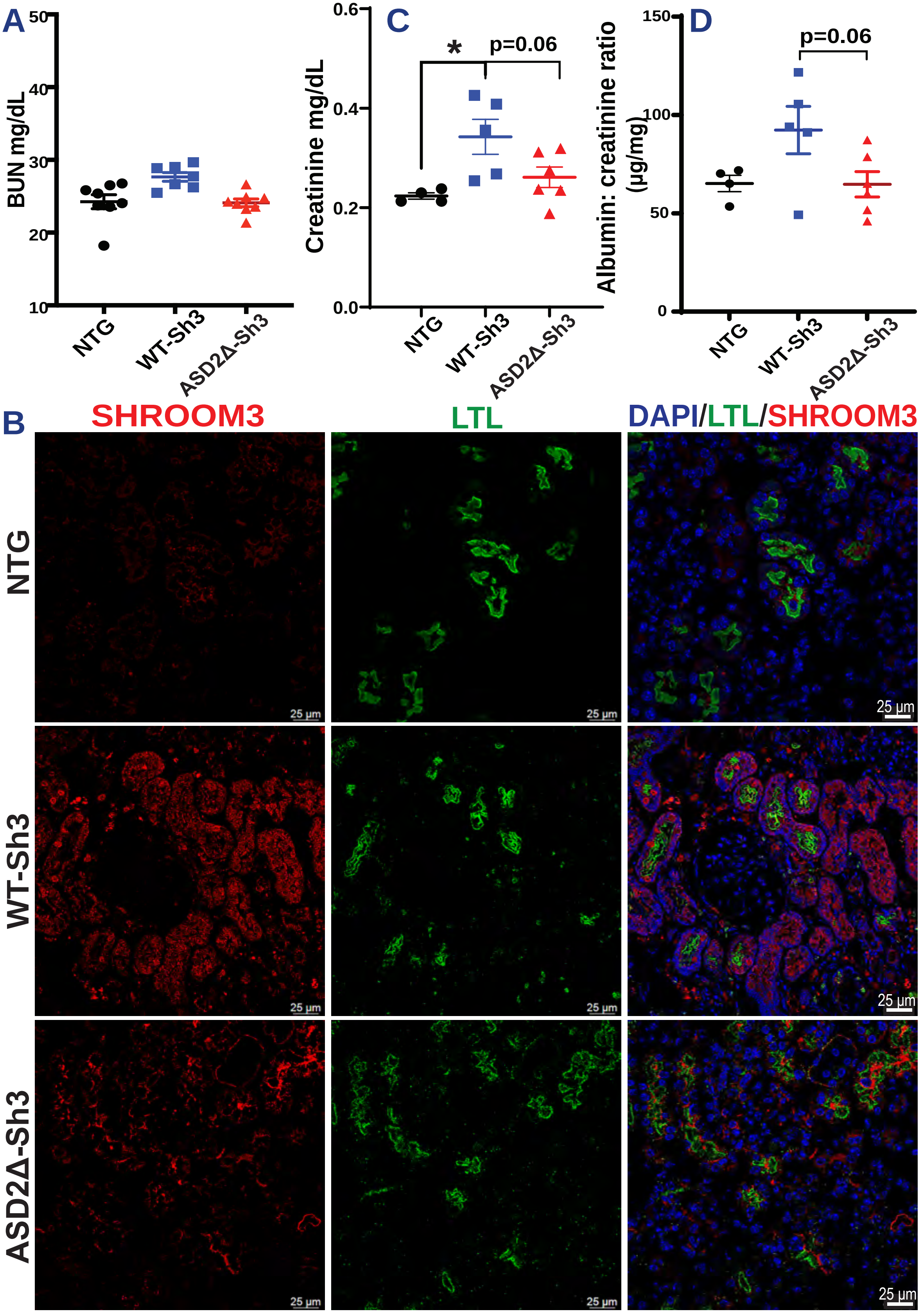
